# Brain Multi-Omic Subtypes of Neuroticism reveal molecular signatures linked to Alzheimer’s Disease

**DOI:** 10.1101/2025.05.19.654904

**Authors:** Andrea R. Zammit, Katia de Pavia Lopes, Caio M.P.F. Batalha, Lei Yu, Victoria N. Poole, Shinya Tasaki, Alifiya Kapasi, Yanling Wang, Philip L. De Jager, Vilas Menon, Nicholas T. Seyfried, Rima Kaddurah-Daouk, Yasser Iturria-Medina, David A. Bennett

**Affiliations:** Rush Alzheimer’s Disease Center, Rush University Medical Center, Chicago, IL, USA; Department of Psychiatry and Behavioral Sciences, Rush University Medical Center, Chicago, IL, USA; Department of Neurological Sciences, Rush University Medical Center, Chicago, IL, USA; Instituto de Assistência Médica ao Servidor Público Estadual, São Paulo, SP, Brazil; Department of Orthopedic Surgery, Rush University Medical Center, Chicago, IL, USA; Department of Pathology, Rush University Medical Center, Chicago, IL, USA; Center for Translational & Computational Neuroimmunology, Department of Neurology, Columbia University Medical Center, New York, NY, USA; Department of Biochemistry and Neurology, Emory University School of Medicine, Atlanta, GA, USA; Department of Psychiatry and Behavioral Sciences, Duke University, Durham, North Carolina, USA; Duke Institute of Brain Sciences, Duke University, Durham, North Carolina, USA; Department of Medicine, Duke University, Durham, North Carolina, USA; Neurology and Neurosurgery Department, Montreal Neurological Institute, Montreal, Canada; McConnell Brain Imaging Centre, Montreal Neurological Institute, Montreal, Canada; Ludmer Centre for Neuroinformatics & Mental Health, Montreal, Canada; McGill University Research Centre for Studies in Aging, Douglas Research Centre.

**Keywords:** brain multi-omics, neuroticism, molecular pseudotime, molecular subtypes, ADRD

## Abstract

**Importance:** Molecular mechanisms linking neuroticism with Alzheimer’s disease traits are unknown.

**Objective:** To identify molecular subtypes of neuroticism and examine their association with ADRD traits.

**Design:** Three ongoing cohort studies were used; Religious Orders Study (ROS), Rush Memory and Aging Project (MAP) and Minority Aging Research Study (MARS), that began enrollment in 1994, 1997, and 2004, respectively.

**Setting:** Older priests, nuns, and brothers from across the U.S. (ROS), older adults (MAP) and older African-American adults (MARS) from across the greater Chicago metropolitan area.

**Participants:** 1,028 decedents with multi-omic data from the dorsolateral prefrontal cortex (DLPFC), the anterior cingulate cortex (AC), and the posterior cingulate gyrus (PCG).

**Exposure(s):** Eight layers of omics (DNA methylation and histone acetylation from DLPFC; RNA seq from AC, DLPFC, and PCG, single-nucleus RNA, TMT proteomics and metabolomics from DLPFC) and Neuroticism using the 12-item version from the NEO Five-Factor Inventory.

**Main outcome(s) and measure(s):** Person-specific multi-omic molecular pseudotime representing molecular progression from low to high phenotypic expression of neuroticism, and three multi-omic brain molecular subtypes of neuroticism which represent distinct omic pathways from no/low neuroticism to high neuroticism that differ by their omic constituents.

Participants are exclusively assigned to the subtype which aligns mostly with their multi-omic profile.

**Results:** The top drivers of subtype differentiation were transcriptomic alterations across three brain regions (DLPFC, AC, PCG) which extensively and differentially characterized the subtypes. The subtypes were also differentially associated with AD pathology, temporal lobe atrophy, and AD dementia, with subtype N_1_ showing the strongest associations.

**Conclusions and Relevance:** Neuroticism may be driven by three distinct molecular subtypes, with subtype N_1_ driving ADRD-related associations, N_2_ showing some ADRD associations, and N_3_ being completely independent of these outcomes. Our data provide novel insights into the biology of individual differences in predispositions of neuroticism and its associations with ADRD traits.

**Key points:** *Question:* What are the brain multi-omics molecular signatures linking neuroticism with Alzheimer’s diseases and related dementias (AD/ADRDs)?

*Findings:* We identified three distinct brain multi-omic molecular subtypes reflecting different molecular pathways underlying neuroticism. Top omic features of the subtypes were extensively and differentially characterized by transcriptomic alterations across three brain regions – dorsolateral prefrontal cortex, anterior cingulate cortex, and posterior cingulate gyrus. Subtype N_1_ was strongly associated with AD pathology, AD dementia, and temporal lobe atrophy.

*Meaning:* The association we typically observe between phenotypic neuroticism and ADRD clinical traits might be largely driven by a molecular pathway underlying this trait.

## Introduction

Neuroticism is the predisposition to experience psychological distress, including anxiety, irritability, anger and negative affectivity.^1^ Because individuals with higher neuroticism are more likely to experience discomfort across situations and over time, it is an indicator of individual differences in chronic psychological distress and is considered a stable trait.^2^ The consequences of having high neuroticism have been widely noted, and include poor health behaviors^3^, adverse mental, physical and cognitive health outcomes^4–8^, and morbidity and mortality.^9^ ^10^ Due to its widespread effects^11^ and its potential modifiability^12^, it is important to understand its neurobiological basis.

We previously showed that neuroticism is associated with cognitive decline, mild cognitive impairment, and Alzheimer’s dementia^13,14^; however, it was not associated with their leading causes including AD pathology, non-AD neurodegenerative pathology and cerebrovascular pathology, and remained associated with ADRD outcomes after adjusting for these pathologies.^15^ Recently, we found that neuroticism was related to select transcriptomic modules^16^, and specific cortical proteins^17^ in the aged prefrontal cortex, which mediated in part the association of neuroticism with cognitive decline, providing evidence that molecular mechanisms are in the pathway to AD dementia.^16^

Here we extend our prior work to uncover the molecular subtypes of neuroticism by unifying, reordering, and stratifying multi-omics patterns using eight layers of omics and an unsupervised multi-modal contrastive trajectory inference algorithm to detect enriched subtypes in a subpopulation in highest tertile of neuroticism relative to a background subpopulation in the lowest tertile of neuroticism.

## Methods

### Study participants

Participants were community-based older adults enrolled in one of three ongoing cohort studies of aging and dementia, the Religious Orders Study(ROS), the Rush Memory and Aging Project(MAP)^18^ or the Minority Aging Research Study(MARS).^19^ Participants are free of known dementia at enrollment, agree to annual clinical evaluation and sign an informed consent and Anatomic Gift Act to donate their brains at death (optional for MARS). All studies were approved by an Institutional Review Board of Rush University Medical Center.^18,19^

### Assessment of Neuroticism

Neuroticism was assessed using the 12-item version from the NEO Five-Factor Inventory administered at or near baseline.^20^ Higher scores (Range0-48) indicate higher neuroticism.

### Multi-omic data origin

We obtained DNA methylation data (N=577) and histone acetylation data (N=494) from the dorsolateral prefrontal cortex (DLPFC); RNA-seq data from the anterior cingulate cortex (AC; N=602), the DLPFC (N=924), and the posterior cingulate gyrus (PCG) (N=564); and single-nucleus RNA-seq data (N=358), TMT proteomics data (N=700) and/or metabolomics data (N=436) from the DLPFC. All data underwent standardized processing and quality-controlled as previously described.^21–24^ Only participants who completed the NEO-FFI at baseline and had at least 2 molecular data modalities were included (N=1208).

### Assessment of postmortem brain neuropathology

Neuropathologic data collection and assessment were performed blinded to all clinical and cognitive data, and followed a standard protocol for tissue preservation, tissue sectioning, and quantification of pathologic findings^25–27^, detailed in **eMethods**. Measured indices of AD/ADRD pathologies included Alzheimer disease, hippocampal sclerosis, TAR DNA-binding protein 43 (TDP-43), neocortical Lewy bodies(LB), macroinfarcts, microinfarcts, cerebral amyloid angiopathy (CAA), atherosclerosis, and arteriolosclerosis.^28^

### Assessment of post-mortem neuroimaging

A subset of 492 participants underwent a postmortem brain imaging protocol, described previously.^29^ Briefly, after one month postmortem, the cerebral hemisphere selected for neuropathological examination was imaged with a multi-echo spin-echo sequence on one of four 3-Tesla MRI scanners, and the resulting images were used for deformation-based morphometry(DBM), detailed in **eMethods**.

### Assessment of cognitive function and cognitive status diagnoses

Detailed annual neuropsychological assessments were administered.^30^ A global composite score was derived by standardizing 19 tests assessing 5 cognitive domains using baseline means and standard deviations across cohorts.^30^ Repeated measures were used to estimate the rate of cognitive decline before death.

AD dementia was diagnosed by an experienced clinician using criteria of the joint working group of the National Institute of Neurologic and Communicative Disorders/Stroke/AD and Related Disorders Association.^31,32^ These criteria require a history of cognitive decline and evidence of impairment in at least two domains of cognitive function, one of which must be memory.^33^

### Assessment of covariates

Date of birth, sex, and years of education were collected at enrollment. Age at death was calculated by subtracting date of birth from date of death.

### Approach

To biologically define participant stratification, we applied an artificial intelligence/machine learning (AI/ML) approach, multi-modal contrastive Trajectory inference (mcTI)^34^. The mcTI algorithm is a novel cross-validated AI/ML algorithm which automatically identifies a series of molecular states (e.g. alteration in DNA methylation, RNA, proteins) reflecting decades of symptom progression and subsequently detects the relative ordering of participants aligned with this progression.^34^ Once we identified molecular subtypes of neuroticism, we statistically analyzed them in relation to their omics composition, their neuropathologic, brain morphologic, and clinical characteristics, outlined below.

#### Unification approach using multi-modal contrastive Trajectory Inference

To identify the brain multi-omics molecular taxonomy underlying predispositions to high neuroticism we unified, reordered, and stratified multi-omics patterns as shown in **Figure 1**. We first stratified participants’ scores on the neuroticism scale into three groups: low(range:0-12), average(range:13-24) and high(range:25-42) as previously described.^16^ This step is required to define a desirable state, referred to as background, and an undesirable state, referred to as target. We defined the background as low neuroticism, and the target as high neuroticism. mcTI then uses the background and target populations to estimate parameters for the model, and subsequently provides two person-specific estimations: **A.)**a person-specific multi-omics molecular progression-score i.e. pseudotime, reflecting the individual’s molecular proximity to an undesirable state (in this case expression of high neuroticism), and **B.)**a putative subtype, corresponding to a distinct molecular subtype in the multi-dimensional omic space of the trait for which participants are exclusively assigned, enabling the identification of the molecular heterogeneity underlying progression. Achieving these two person-specific estimations, consists of six steps as previously detailed^34^ and described in the **eMethods**.

**Figure 1.**
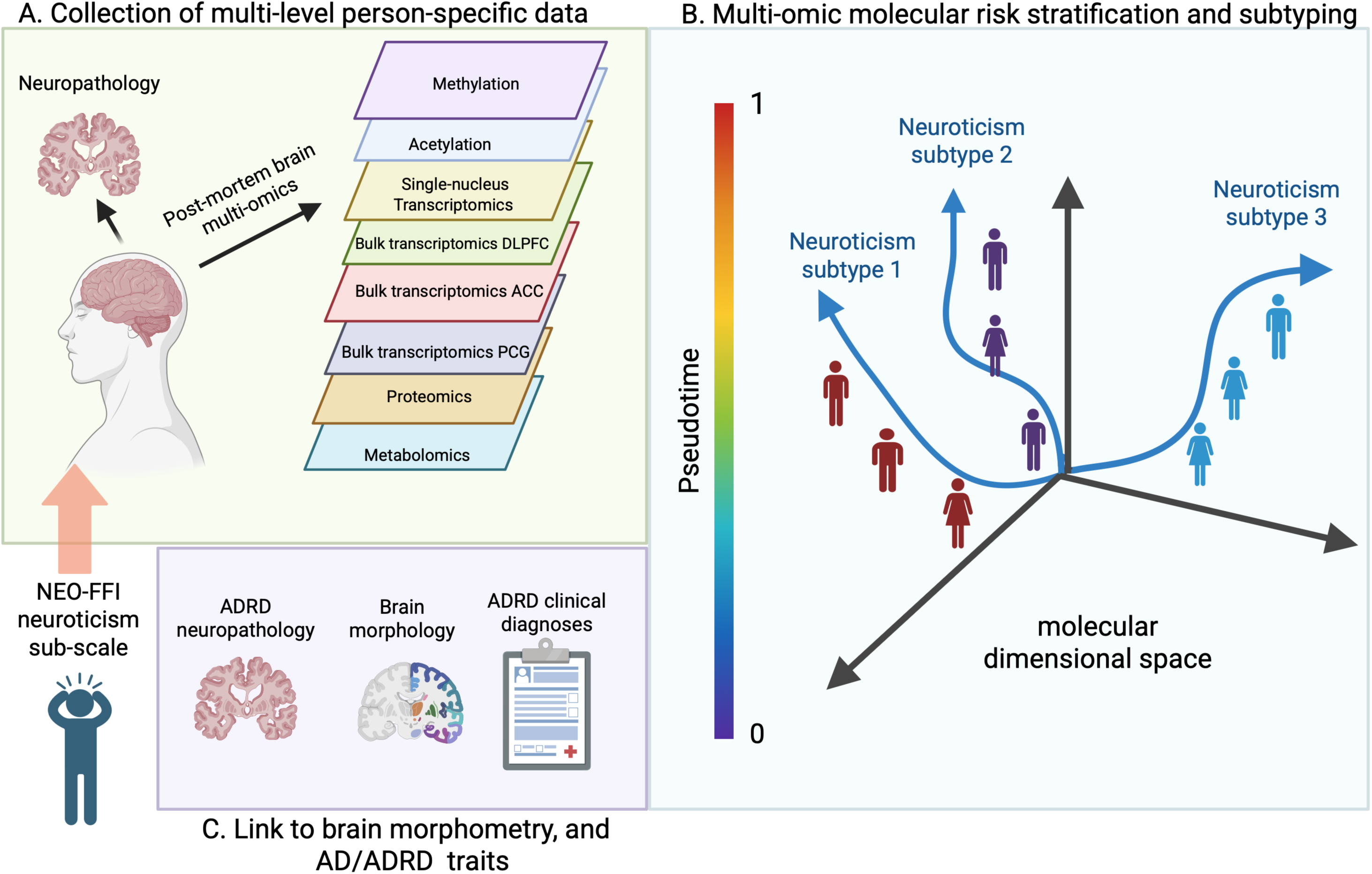
Schematic of study design. **Panel A.** We included multi-modal antemortem and postmortem data from participants who completed the NEO-Five Factor Inventory neuroticism sub-scale at study baseline and who had at least two omic-modalities from eight layers of post-mortem brain multi-omic molecular data (N=1,208). **Panel B.** We applied a contrastive artificial intelligence/machine learning algorithm to integrate the brain’s multi-omics molecular data, to obtain a multidimensional phenotypic space where participants define biologically distinctive subtypes from no/low neuroticism to high neuroticism. For each participant we obtained a molecular progressive index, pseudotime score, and subsequently three neuroticism subtypes. **Panel C.** We then tested whether the brain-derived pseudotime and subtypes predict antemortem ADRD clinical traits, and postmortem ADRD pathological traits and brain morphometric indices.

### Characterization of the pseudotime and identified subtypes

We characterized the top features contributing to the pseudotime calculation and subtypes. We used chi-squared tests to determine whether observed omic modalities differed by subtype.

### Associations with ADRD pathologic traits

We compared frequency of pathologies across subtypes using chi-square tests. We tested associations between pseudotime and three continuous ADRD neuropathologic indices i.e., global AD pathology, amyloid-β, and PHF-tau in separate linear regressions. We ran logistic regression models for the following binary outcomes: pathologic diagnosis of AD, presence of neocortical LBs, hippocampal sclerosis, TDP-43 cytoplasmic inclusion extended beyond amygdala, presence of macroscopic infarcts, microinfarcts, moderate/severe arteriolosclerosis, and atherosclerosis. Similar analyses tested whether the subtypes are associated with these pathologic indices.

### Associations with neuroimaging

We tested associations with brain morphometry by running two voxel-wise general linear models of deformation via FSL PALM^35,36^ which accounted for differences in both mean and variance across scanners. Pseudotime served as the explanatory variable of interest for the first model, while indicator variables representing the background and subtypes were of interest for the second model. To compare deformation across groups in the latter model, we contrasted effect maps for each group pair. Both models were adjusted for demographics, postmortem interval, and scanner. P-values were computed^37^ and the threshold-free cluster enhancement approach was used to define clusters of significance. Associations were considered statistically significant at p≤.05, family-wise error-rate(FWER) corrected.

### Associations with ADRD clinical traits

#### Pseudotime

We tested the association of pseudotime with cognitive decline by using a linear mixed effects model. The model included terms for age at death, sex, education, pseudotime, and time in years from death, and the interaction of each measure with time before death. The term for pseudotime estimates the association of pseudotime with the level of cognition at death, and the interaction estimates the association of pseudotime with annual rate of cognitive decline. We tested the association between pseudotime and dementia using a logistic regression model.

#### Subtypes

We ran linear-mixed effects models by including the subtypes as the binary predictors adjusting for the same covariates. We modeled the background as the reference. The term for each subtype estimates the difference in the level of cognition between the subtype and the background, and the interaction estimates the difference in the annual rate of cognitive decline between the subtype and the background. We then re-ran the same models by further adjusting for the 9 ADRD neuropathologic indices. We ran logistic regression models to test whether the subtypes are associated with dementia.

## Results

A total of 1028 participants from MAP(n=497), ROS(n=520) or MARS(n=11) were included. Mean age at study baseline was just over 80 years and mean age at death was close to 90 years. Almost two-thirds were female, and most participants completed college. Mean score on neuroticism was 16.6(SD=6.4)(**Table 1**).

**Table 1.**
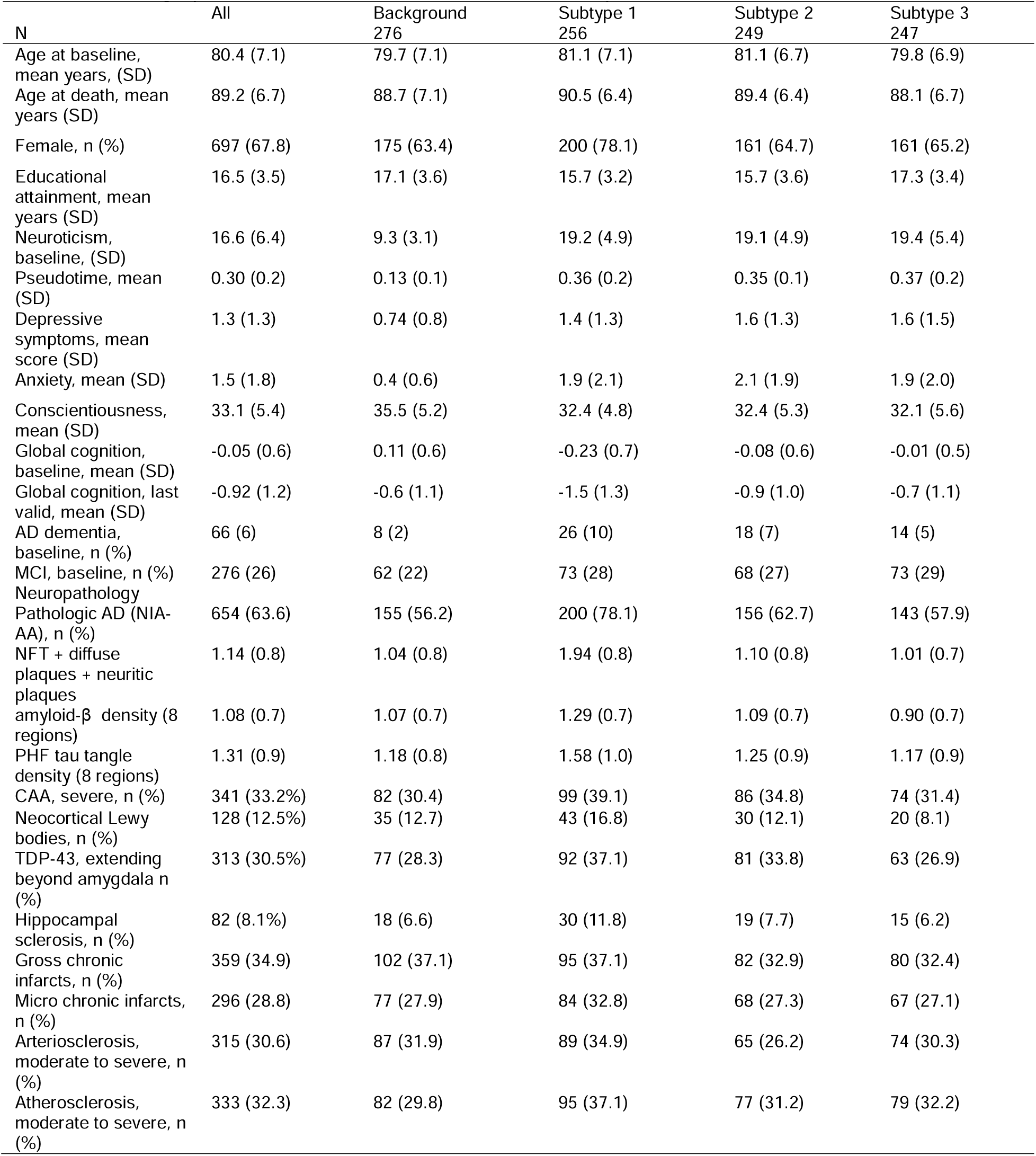
Demographic and clinical characteristics of the sample.

### Characterizing the multi-omic brain molecular pseudotime and distinct molecular subtypes of neuroticism

Pseudotime was mostly informed by transcriptomic alterations in the PCG and the AC (**Figure 2A**). Other features included metabolomic and epigenomic (mainly acetylation) alterations. The top contributor was the Mediterranean Fever(*MEFV*) gene, which encodes a protein called pyrin, and is involved in the body’s immune response particularly in regulating inflammation. Overactivation of this gene can lead to an overactive immune system.

**Figure 2.**
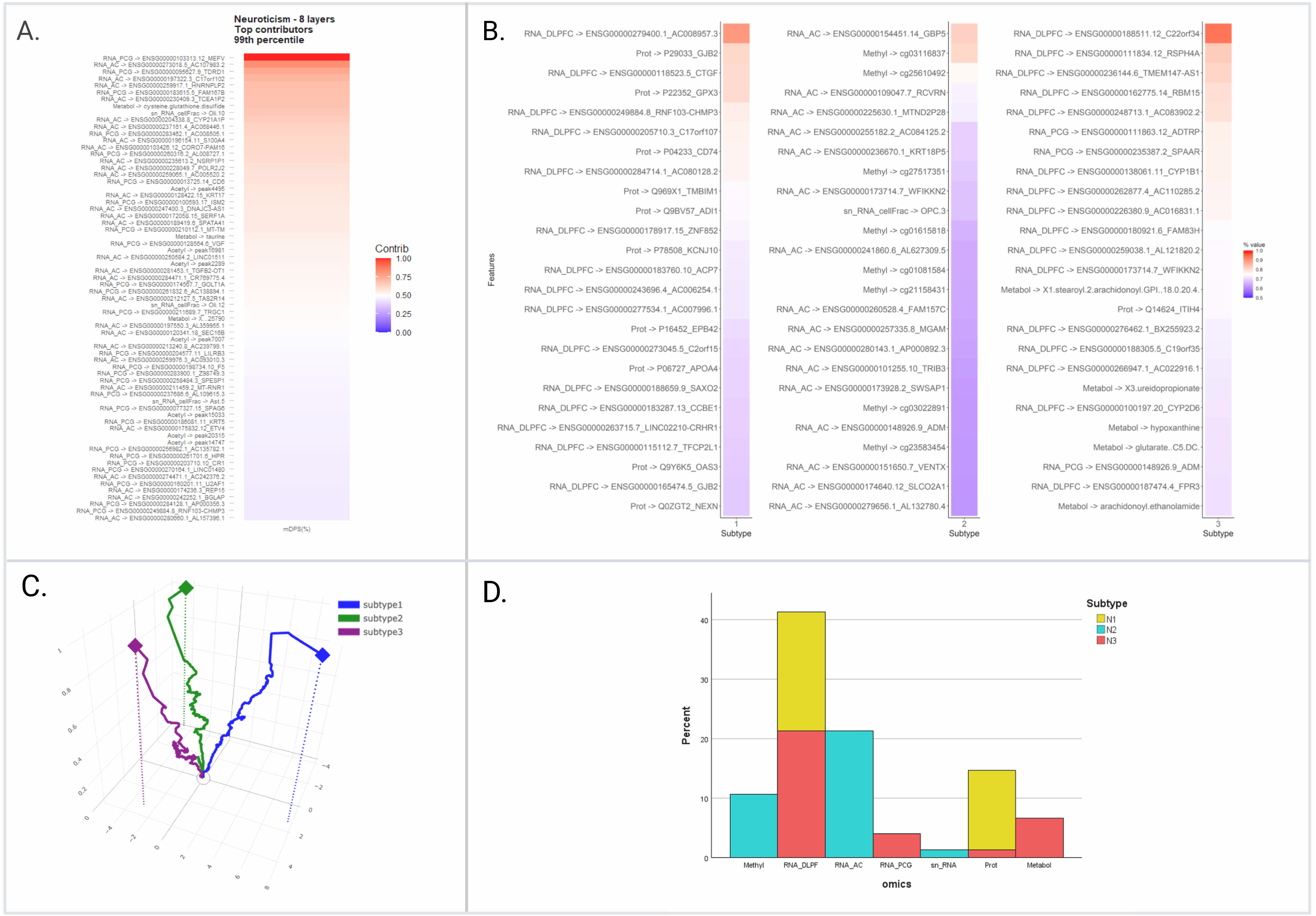
A. Characterization of the top 1% of omic markers of the neuroticism pseudotime. **B.** Top 25 omic markers of the three subtypes. **C.** 3-D characterization of the three identified subtypes. **C.** Distribution of omic markers per subtype. **D.** Bar plot showing a color-coded distribution of top omic features per subtype.

Brain multi-omic molecular information revealed three distinct subtypes underlying neuroticism which we will refer to as N_1_, N_2_, and N_3_. Here, the algorithm detected three subgroups of participants which corresponded to a distinct concatenation of participants in the integrated molecular space.

Transcriptomic alterations in three brain regions extensively and differentially characterized the subtypes. While 60% of the top 25 omics features characterizing N_1_ were RNA alterations in the DLPFC, 64% of the top features characterizing N_2_ were identified in the AC and a further 4% in single-nucleus RNA. Meanwhile RNA alterations in N_3_ were from both the DLFPC (64%) and the PCG (12%). **Figure 2B** shows the top 25 influential omics features per subtype; **Figure 2C** shows these subtypes in 3-dimsenional molecular space; **Figure 2D** shows the distributions of the top omic features across subtypes.

### AD/ADRD neuropathology across the subtypes

The frequency (range:0-9) and combination of pathologies were very heterogeneous both within and across groups (**S**.**Figures 1–4).** However, N_1_ had the highest mean number of pathologies (3.3, SD=1.6) relative to background and all subtypes (p<0.001; **S.Table 1**).

N_1_ had more pathologic AD (78%) than background and the subtypes Over 82% of participants in N_1_ had pathologic AD and/or CAA (χ^2^(3, 1007)=27.3, p<0.001). Almost half had non-AD neurodegenerative pathology (χ^2^(3, 988)=14.2, *p*=0.003); however, there were no differences between N_1_ and N_2._ A third of all participants with cortical LBs were in N_1_. There were no differences for frequency of vessel pathology (χ^2^(3, 1015)=5.01,p=0.171), or cerebral infarcts (χ^2^(3, 1028)=2.67, *p*=0.445) (**Figure 3 panels A-D**).

**Figure 3.**
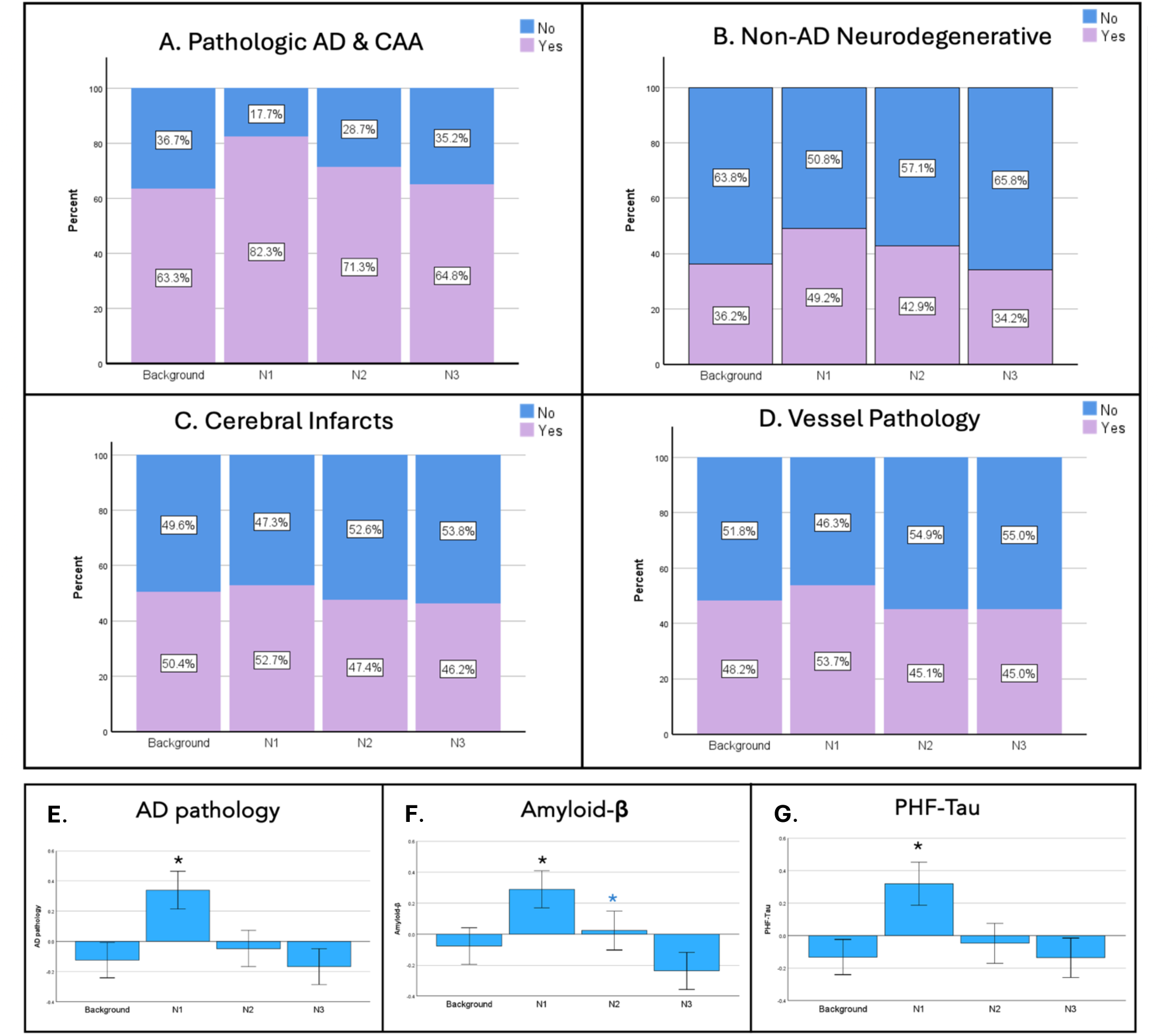
Percentage of individuals with ADRD pathologic indices stratified by background and three multi-omic molecular neuroticism subtypes. Panels A-D. CAA and pathologic AD includes presence of moderate to severe presence of cerebral amyloid angiopathy and an NIA-Reagan diagnosis of AD. Non-AD neurodegenerative pathology includes neocortical Lewy bodies, TDP-43, and hippocampal sclerosis). Cerebral infarcts include gross and/or micro infarcts. Cerebral vessel pathology includes arteriosclerosis and atherosclerosis. **Panels E-G.** Bar charts illustrating the differential association of the Neuroticism subtypes with a summary measure of AD pathology, and its constituents, amyloid-β and PHF-tau tangles. Bonferroni-adjusted post-hoc analyses showed that N_1_ had significantly more AD pathology (p<0.001), amyloid-β (p<0.001) and PHF-tau (p<0.001) than Background, N_2_ and N_3_. The black asterisks show these associations. N_2_ had significantly more amyloid-β than N_3_. (*p*=0.02). The blue asterisk shows this association. The measures are z-transformed for illustration purposes. The means are adjusted for age, sex, and education.

**Figure 4.**
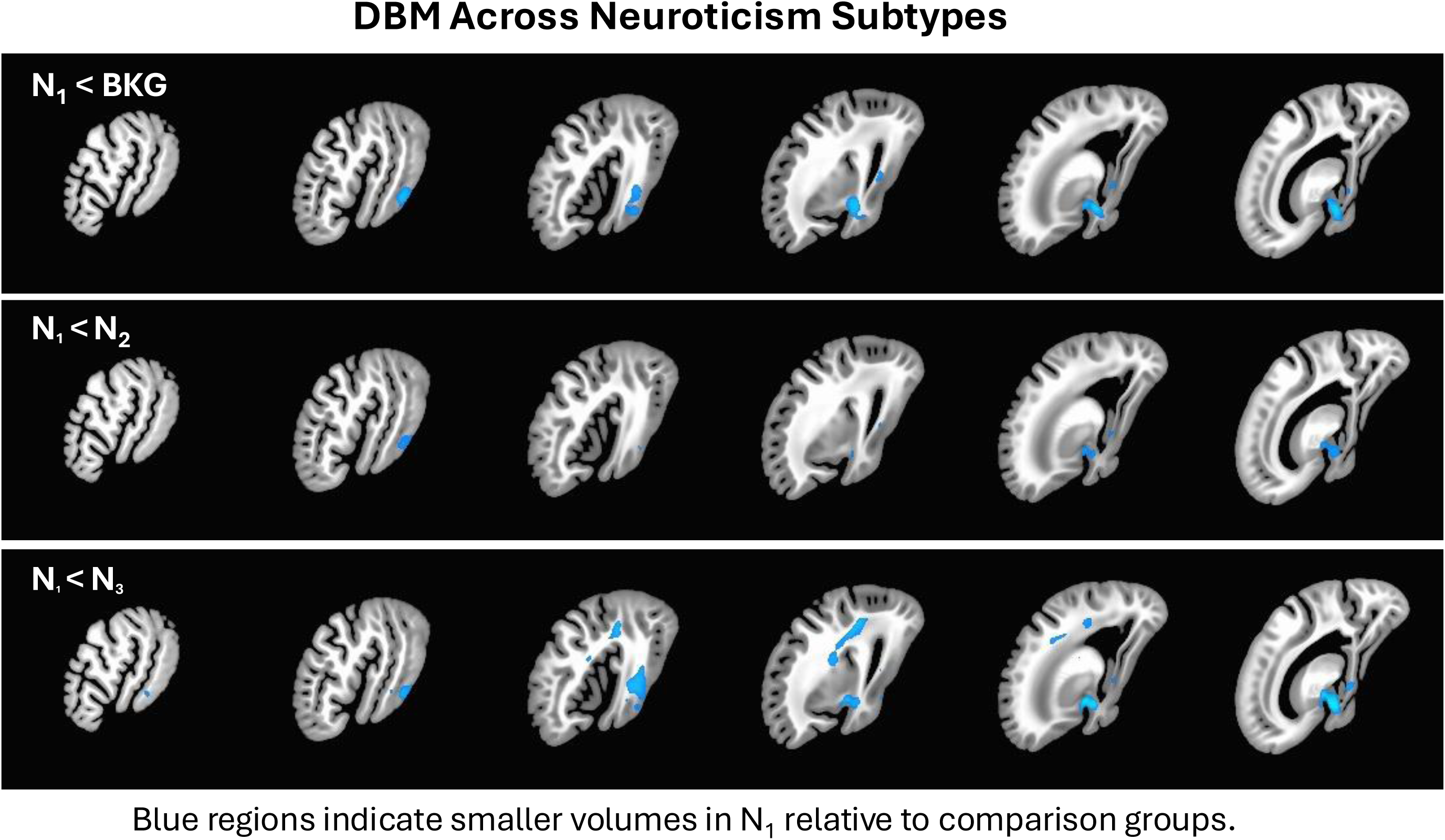
Background and inter-subtype differences in brain morphology (FWER *p***≤**.05). Relative to background, N_1_ had smaller regions in the dorsal and medial temporal lobes. Relative to N_2_, N_1_ had smaller regions in the lateral inferior temporal gyrus, the medial temporal lobes including amygdala and anterior hippocampus, and neighboring white matter. Relative to N_3,_ N_1_ had smaller regions in the temporal and medial temporal lobe, as well as in the parietal white matter.

Continuous AD neuropathologic indices indicated mean differences on global AD pathology Post-hoc analyses showed that N_1_ had significantly more AD pathology, more amyloid-β, and more PHF-tau than the background or any of the subtypes (**Table 1**). N_2_ also had significantly more amyloid-β relative to N_3_ (**Figure 3, panels E-G)**.

Pseudotime was not associated with any neuropathologic indices indicating that this molecular index is not driven by pathology in neuroticism.

### Post-mortem brain morphology across the subtypes

We examined differences in brain morphometry across a subset of 492 participants (background=142; N_1_=135; N_2_=114; N_3_=101) with postmortem MRI. We observed that N_1_ had smaller temporal and medial temporal volumes than the Background, N_2_, and N_3_ (**Figure 4**). Relative to background, N_1_ exhibited smaller dorsal and medial temporal lobe regions. Relative to N_2_, N_1_ had smaller regions in the medial temporal lobe, specifically the amygdala and the anterior hippocampus, as well as in the lateral inferior temporal gyrus and neighboring white matter areas. Relative to N_3_, N_1_ had smaller regions in the vicinity of the temporal lobe, and in the white matter regions of the parietal lobe.

We did not observe any other differences, nor did we observe deformation associations with pseudotime.

### AD/ADRD clinical outcomes across the subtypes

Higher pseudotime was associated with a faster rate of cognitive decline (est.=-0.052, SE=0.02, p=0.02) as expected given the association of neuroticism with cognitive decline. **S**.Figure 5 shows that compared to those with average pseudotime (50^th^percentile), decline was 12% faster for those in the 90^th^percentile, and 9% slower for those in the 10^th^percentile. Higher pseudotime was associated with higher odds of dementia proximate to death (OR=2.38, 95%CI=1.14-4.96, p=0.02).

Relative to background, N_1_ had faster cognitive decline before death (est.=-0.056, SE=0.01, *p* <0.001); however, decline in N_2_ was borderline significant while N_3_ was not significantly differsent (N_2_: est.=-0.019, SE=0.01, p=0.05; N_3_: est.=0.006, SE=0.01, p=0.571). The association of N_1_ with cognitive decline persisted but was attenuated after we adjusted for 9 neuropathologic indices (N_1_ est.=-0.029 SE=0.01, p=<0.001). By contrast, the association of N_2_ with cognitive decline was essentially unchanged (N_2_ est.=-0.017, SE=0.008, p=0.039).

Odds of dementia was over 3 times higher in N_1_ than in background (OR=3.34, 95%CI=2.31-4.83, p<0.001); borderline significant in N_2_ (OR=1.45, 95%CI=1.01-2.09, p=0.05), and not significant in N_3_ (OR=0.98, 95%CI=0.68-1.43, p=0.93).

## Discussion

We investigated the brain molecular multi-omic taxonomy underlying the phenotypic expression of neuroticism using a unified artificial intelligence/machine learning framework. By integrating eight layers of omics across three brain regions in over 1,000 individuals we identified a person-specific molecular progression-index, pseudotime, indicating molecular proximity to high expression of neuroticism. Decomposition of pseudotime revealed three distinct molecular pathways underlying individual differences in predisposition to neuroticism. Transcriptomic alterations across three brain regions extensively and differentially characterized the subtypes. Transcriptomic alterations in N_1_ were primarily detected in the DLPFC, whereas N_2_ was marked by differences in the AC, and N_3_ showed alterations in both PCG and DLPFC. Notably, N_1_ had strong associations with AD pathology, brain atrophy, AD dementia, and cognitive decline before and after regressing out effects of pathology, indicating that the robust associations we typically observe between neuroticism and ADRD related outcomes is largely driven by this molecular pathway. While some of these associations were also observed in N_2_ they were clearly weaker, and they were absent in N_3_. Together, these results identify a relatively specific molecular pathway linking neuroticism with ADRD pathologic and clinical traits. These differential effects were supported with structural imaging morphometric differences by subtype.

Previous studies have shown that chronic psychological distress profoundly impacts that brain’s structural^11,38^ and molecular^39,40^ makeup. We also previously identified associations between neuroticism and 18 cortical co-expressed modules in the aging prefrontal cortex ^16^ as well as associations with specific cortical proteins.^17^ A subset of four modules^16^ and two proteins^17^ were also linked to cognitive decline and resilience, suggesting molecular pathways by which neuroticism may influence ADRD outcomes. However, these studies were limited to single-layer omics. More recently, we identified three multi-omic brain molecular subtypes of AD dementia, each reflecting a unique molecular trajectory from no cognitive impairment to AD dementia.^34^ We subsequently found that these three AD subtypes were differentially associated with neuroticism providing evidence that neuroticism may contribute to ADRD risk via shared multi-omic mechanisms.^41^

We extend prior work by directly characterizing the multi-omic molecular signatures of neuroticism and examining their relevance to ADRD outcomes. To our knowledge, this is the first study to integrate multiple omics layers to investigate the neurobiology of neuroticism.

Subtype N_1_’s top contributing feature was AC008957.3 which is a long non-coding RNA, associated with the *SLC1A3-AS1* gene on chromosome 5. This chromosome contains clusters of genes involved in immune signaling, including interleukins^42^ though it has not been previously linked to neuroticism.^43^ *SLC1A3-AS1* is an antisense gene to *SLC1A3,* which encodes a glutamate transporter critical for terminating excitatory neurotransmission in the CNS. While the specific function of AC008957.3 is still unknown, it may regulate *ALC1A3* activity. N_2_^’^s top feature was *Guanylate Binding Protein 5 (GBP5)*, a protein-coding gene on chromosome 1, involved in immune response and inflammation. Although chromosome 1 has been implicated in neuroticism^39^ *GBP5* has not been previously linked to this trait. Finally, N_3_’s top contributor, was *C22orf34* a long non-coding RNA on chromosome 22. While the functions of *C22orf34* is unclear, a nearby gene (*C22orf36*) has been associated with neuroticism in previous GWAS studies^44^, and *C22orf34* has been linked to immune response in diffuse large B-cell lymphoma.^45^ Collectively, our findings suggest that immune response and inflammation are key components of the molecular heterogeneity underlying neuroticism, consistent with prior work.^46^

Our findings suggest that the impact of neuroticism on ADRD outcomes is not uniform across individuals. Two individuals with similar levels of neuroticism may have vastly different molecular profiles, and consequently distinct health risks. Previous genome-wide studies identified genetically distinguishable subclusters of neuroticism, pointing to divergent causal mechanisms.^44,47,48^ We also previously identified four co-expressed gene modules differentially associated with neuroticism, cognitive decline and amyloid-ß and tau pathology suggesting multiple mechanistic pathways through which neuroticism may impact cognitive decline.^16^

Future work needs to clarify the sequence of molecular events triggered or modified by chronic predispositions to neuroticism. For instance, the ADRD associations observed in N_1_ may reflect downstream consequences of a sequalae of molecular alterations triggered by persistent levels of neuroticism. Alternatively, neuroticism may interact with ongoing neuropathological processes to exacerbate risk. Our prior work suggests that the association of neuroticism with cognitive decline is more likely mediated by molecular pathways impacting downstream pathology accumulation, than through a direct association^16^, though more work is needed to establish directionality.

These findings have clinical implications. Neuroticism is strongly related to multiple psychiatric conditions, including depression,^44,49^ with growing evidence for shared biological pathways across these traits.^50^ Future work will need to elucidate common molecular underpinnings with depressive symptoms and anxiety. Targeting these shared pathways with pharmacological agents may “open up windows” of neural plasticity^11,38^, enhancing the effectiveness of cognitive and behavioral interventions, potentially altering cognitive aging trajectories and mitigating later-life cognitive impairment.

This study has limitations and strengths. Neuroticism is a complex trait, and the brief measure used in this study precluded the adequate investigation of all facets included in the 48-item scale. However, we previously reported a .90 correlation with shorter-item versions of the NEO^51^ testifying to their validity. We have few minoritized groups; 99% of participants were from ROS or MAP, who are mostly non-Hispanic white. Future work should replicate these findings in diverse populations. However, rates of participation in the clinical evaluations and brain autopsy in the ROS and MAP cohorts are high, minimizing bias due to selective attrition.

## Supporting information

eMethods

## Data sharing

The data are available via the Rush Alzheimer’s Disease Center Research Resource Sharing Hub. Qualified applicants should complete an application including study premises and a brief description of the research plan.

Brain multi-omic data are available in the AMP-AD knowledge portal (www.synapse.org), using the following Synapse IDs: syn3157275 (epigenomic data), syn3388564 (transcriptomic), syn23650894 (single cell RNA seq), syn17015098 (TMT proteomic), and syn26007830 (metabolomic). The metabolomic data were generated by the Alzheimer’s Disease Metabolomics Consortium (ADMC).

## Acknowledgements

We appreciate the participants of MAP for the time generously given for data collection and for consenting for brain donation. We also acknowledge staff of Rush Alzheimer’s Disease Center for data collection, management, and analyses.

This work was supported by National Institutes of Health: R01AG17917, P30AG10161, P30AG72975, R01AG015819, U01AG61356, U01NS100599, R01AG064233, RF1NS139975, P30 AG072975. We would also like to thank Michael Urbut and the Paul M. Angell Family Foundation for their generous financial support given towards this work.

The funding organizations had no role in the design or conduct of the study; collection, management, analysis, or interpretation of the data; or preparation, review, or approval of the manuscript.

