## Supplementary material for "Brain Multi-Omic Subtypes of Neuroticism reveal molecular signatures linked to Alzheimer’s Disease": eMethods

**eMethods for**

**Unified multi-omic molecular taxonomy of neuroticism**

**Zammit A.R. et al.**

**Assessment of Neuroticism**

Neuroticism is an indicator of proneness to psychological distress. Participants rated agreement with each item on a 5-point Likert scale ranging from 0 (strongly disagree) to 4 (strongly agree). The total score in participants with the 12-item scale was computed by summing item scores to yield a total score that ranged from 0 to 48, with higher scores indicating more proneness to psychological distress. Internal consistency reliability as determined by the Cronbach coefficient alpha was 0.8, which indicates adequate internal consistency.

**Assessment of postmortem brain neuropathology**

The cerebral hemispheres were coronally cut into 1-cm slabs, and the brainstem was removed at the level of the mamillary bodies and bisected at the mid pons level. One hemisphere was prepared for histologic evaluation and the other hemisphere was frozen for the collection of omics. The fresh slabs were fixed in 4% paraformaldehyde. Tissue blocks from predetermined regions were dissected, embedded in paraffin, and cut into 6 and 20 microsections^1^.

**Assessment of post-mortem neuroimaging**

The parameters of the ex-vivo MRI protocol were similar for the four MRI scanners used in this work, and have been presented in detail in reference 50. The ex-vivo brain MRI images were masked, corrected for field inhomogeneity using the N4 approach^2^, and were non-linearly registered to an in-house postmortem hemispheric template^3^ using Advanced Normalization Tools (ANTs)^4^. The determinant of the Jacobian of the transformation was calculated at the voxel-wise level, and the resulting maps were smoothed with a 4mm FWHM Gaussian kernel. Higher values in these smoothed maps indicated the presence of more tissue that had to be contracted through image registration to fit the template, whereas lower values indicated less tissue that had to be expanded to fit the template.

**Steps in conducting** **multi-omics contrastive Trajectory Inference as outlined previously^5^**

i) Data adjustment. Data adjustment included adjustment for age, sex, and education.
ii) Contrastive principal components analyses (cPCA). Data exploration and visualization were performed using cPCA^6^ to reduce each data modality to a few components capturing enriched patterns of the neuroticism trait. cPCA allows each participant to be represented in a reduced n-dimensional space where the corresponding position reflects (higher or lower) phenotypic expression. This technique identifies low-dimensional patterns that are enriched in a target dataset (here, high neuroticism) relative to a comparison background dataset (low neuroticism). By controlling for the effects of characteristic patterns in the background, cPCA allows the visualization of specific data structures missed by standard data exploration and visualization (such as standard PCA). Specifically, if C_target_ and C_background_ are the covariance matrices of the target and background data, the direction returned by cPCA are the singular vectors of the weighted difference of the covariance matrices: C_target_ - α.C_background_. The contrast parameter α represents the trade-off between having the high target variance and the low background variance. Multiple values of α are used (i.e. 10 logarithmically equally spaced points between 10^-2^ and 10^2^). Instead of choosing a single α, the resulting subspaces for all the values are clustered based on their proximity in terms of the principal angle and spectral clustering^7^ in a few subspaces. The data are then projected onto each of these few subspaces, revealing different trends within the target data. The original cPCA algorithm selects the final subspace via visual examination, however we automatically select the subspace that maximizes the clustering tendency in the target population.
iii) Probabilistic principal components analysis (pPCA). The contrast principal components from all the data modalities were subsequently used as input into a probabilistic Principal Components Analysis (pPCA); this analysis identified fused contrasted components capturing the common variance across all modalities’ enriched patterns, while dealing with missing data across the different omics layers.
iv) Subject-subject dissimilarity network (SDN). Based on the inter-individual Euclidian distances across the resulting fused PCs we then ran a subject-subject dissimilarity network. This SDN is used to calculate the shortest path from any participant to background group. The concatenation of relatively similar subjects in the integrate multi-omics molecular space that minimizes the distance to the background’s centroid is defined as shortest path. The position of each subject in the corresponding shortest path reflects the individual distance to the background’s population (here, low neuroticism). To quantify the distance from low to high neuroticism, the person-specific multi-omics pseudotime is calculated as the shortest distance value to the background’s centroid, relative to the maximum population value. Values are standardized thus ranging from 0 (closer to background) to 1 (closer to target).
v) T-SNE. The fused contrasted components are statistically adjusted by the pseudotime values via robust regression. These now adjusted components are then non-linearly embedded into a two- or three- dimensional space via t-SNE^8^, where participants are subsequently clustered. The number of t-SNE dimensions is selected to maximize results stability across perplexity values (i.e. in the range [5 to 50] with 5 as step size) and clustering criteria. The resulting clusters then define then final putative subtypes. Optimal number of clusters/subtypes are defined by using a majority rule across the Calinski-Harabasz and Silhouette criteria.^9,10^ The subtypes’ stability and significance are evaluated via randomized permutations. Subtype stability is defined as the rate at which pairs of subjects group together into the same subtypes upon repeated clustering on random subsets of the input data.^11^ Analyses were performed in MATLAB version R2021b.
vi) Distinct subtype contributors. The objective here was to identify distinct molecular alterations across each subtype by detecting omic features that significantly deviate from typical values in the background group (p<0.05, FDR-corrected), while remaining statistically stable in other subtypes. An ANOVA test with 500 randomized permutations was run for each subtype *i* and omic feature *j*. This was done to calculate the alteration levels of each subtype, thus generating F_ij_ and P_ij_ statistics, adjusted for age, sex, and education. All omic features that were significantly different for any subtypes were identified (p<0.05, FDR-corrected). From this pool we then selected the omic features that were significantly altered for subtype *i* but not significantly altered for the other subtypes. These selected omic features constituted the distinct molecular signature for subtype *i.* the selected features within subtype *i*, were then also ranked according to the propensity to be predominantly different from *i* while being preserved in the other subtypes. To quantify this tendency, we calculated an “anisotropy” index, defined as follows: Ai,j = $\frac{{(Fi,j-\hat{F_{j}})}^{2}+{(Fk,j-\hat{F_{j}})}^{2}+{(Fm,j-\hat{F_{j}})}^{2}}{{Fi,j}^{2}+{Fk,j}^{2}+{Fm,j}^{2}}$ , where k and m represent the other subtypes, and $\hat{F_{j}}$is the average F-value for feature j across subtypes. This formula can accommodate different numbers of subtypes by including or excluding terms accordingly. The index (*Aij*) yields a scaler value between zero and one, indicating the distinction of feature *j* concerning subtype *i*; values closer to one suggest stronger distinction. This approach is an extension of the concept of fractional anisotropy commonly employed in diffusion neuroimaging^12^ and relates to eccentricity of conic section in three dimensions, normalized to a unit range. Notice that while F-values may not be directly comparable across data modalities due to varying sample sizes, the *A_i,j_*_,_ values allow for meaningful comparisons since they are standardized across subtypes.

**Assessing markers contributions on pseudotime.** For each omics modality, the total contribution C_i_ of each modality-specific markers *i* to the obtained reduced representation space (and the multi-omics pseudotime) was quantified as

$$C_{i}= 100\cdot\sum_{k=1}^{N_{pPCAc}} \lambda_{k}^{pPCAc,norm}\cdot\sum_{j=1}^{N_{cPCAc}} \left( \lambda_{j}^{cPCA, norm}\cdot\frac{\omega_{i,j}^{2}}{\sum_{k=1}^{N_{features}} \omega_{i,j}^{2}} \right)$$

where $\lambda_{k}^{pPCAc,norm}$ is the normalized eigenvalue of the pPCA component *k* from a total of $N_{pPCAc}$ resulting components after fusing all data modalities, $N_{cPCAc}$ is the number of contrasted principal components for marker *i*’s corresponding data modality, $\lambda_{j}^{cPCA,norm}=\left( \lambda_{j}-\min\_\lambda\right)/{max\left( \lambda_{m}-\min\_\lambda\right)}$ is the normalized eigenvalue of the contrasted principal component *j*, $\min\_\lambda$ is the minimum obtained eigenvalue across all markers for *j*, $\omega_{i,j}$ is the loading/weight of the marker *i* on the component *j*, and $N_{markers}$is the total number of modality-specific markers in *i*’s corresponding data modality.

**Assessing omics contributions on subtyping.** For each omic modality *i*, its contribution to the obtained subtypes was calculated as:

i) *i* was removed from the *mcTI* algorithm’s input data, and a new set of subtypes was obtained (i.e., without any information from *i*),
ii) an index reflecting the level of dependence in modality *i* for obtaining the original subtypes: $1-nMI$, where $nMI$ is the *normalized Mutual Information* between the two obtained subtyping configurations (i.e., with and without considering modality *i*). Notice that a value of 1 would imply a high level of dependency in modality *i*, and 0 otherwise.
