## Supplementary Material for "Brain Multi-Omic Subtypes of Neuroticism reveal molecular signatures linked to Alzheimer’s Disease"

**Unified multi-omic molecular taxonomy of neuroticism**

**Zammit AR et al.**

**eTable 1.** Number of pathologies stratified by background and neuroticism subtypes.

| Frequency of pathologies | Background | Neuroticism subtype 1 | Neuroticism subtype 2 | Neuroticism subtype 3 |
| --- | --- | --- | --- | --- |
|  | N (%) | N (%) | N (%) | N (%) |
| 0 | 20 (7.6) | 9 (3.6) | 13 (5.5) | 26 (11.6) |
| 1 | 60 (22.7) | 28 (11.3) | 47 (20.0) | 36 (16.0) |
| 2 | 56 (21.2) | 34 (13.8) | 52 (22.1) | 65 (28.9) |
| 3 | 48 (18.2) | 77 (31.2) | 51 (21.7) | 43 (19.1) |
| 4 | 47 (17.8) | 50 (20.2) | 44 (18.7) | 22 (9.8) |
| 5 | 20 (7.6) | 31 (12.6) | 21 (8.9) | 22 (9.8) |
| 6 | 12 (4.5) | 12 (4.9) | 5 (2.1) | 9 (4.0) |
| 7 | 1 (0.4) | 6 (2.4) | 2 (0.9) | 1 (0.4) |
| 8 | 0 | 0 | 0 | 0 |
| 9 | 0 | 0 | 0 | 1 (0.4) |

**Figures**

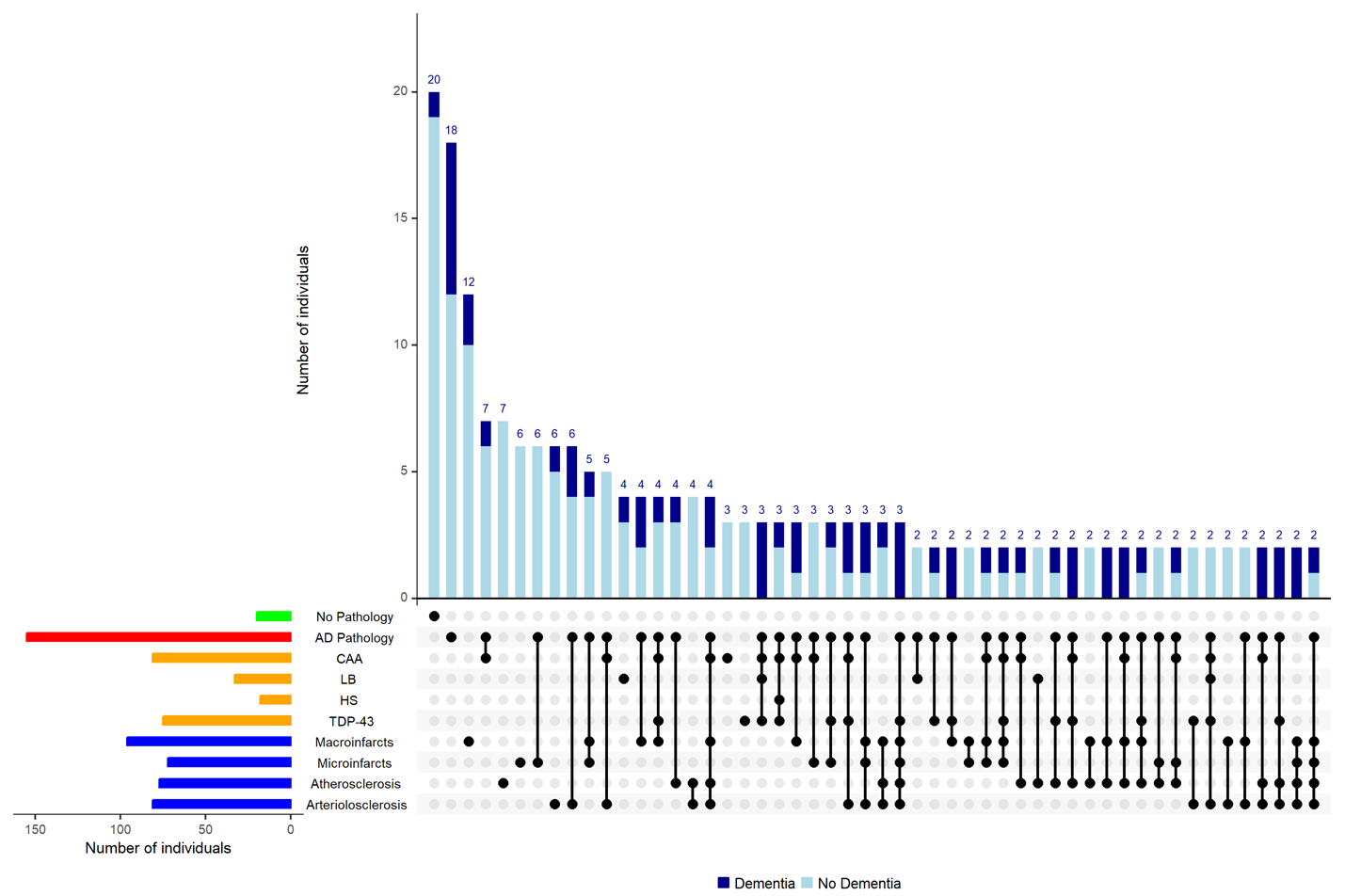

**S. Figure 1. UpSet figure of the distribution and frequency of mixed pathologies for background.** The bar chart in the lower left corner shows the frequencies of individual neuropathologic indices investigated in this study. Connected black dots on the x-axis indicate the specific combination of neuropathology represented (we show only combinations of 2+ pathologies). Histograms in the main panel show the frequencies of the neuropathologic indices for participants with and without dementia proximate to death, ordered by their frequency. The height of each bar corresponds to the number of persons with each combination. As illustrated in the figure, neuropathologic indices frequently co-occur and the most common at pathologic AD, followed by pathologic AD and a neurodegenerative pathology, most commonly, cerebral amyloid angiopathy. AD=Alzheimer’s disease. CAA= cerebral amyloid angiopathy; LB=Lewy bodies. HS=hippocampal sclerosis. TDP-43=transactivate response DNA-binding protein 43.

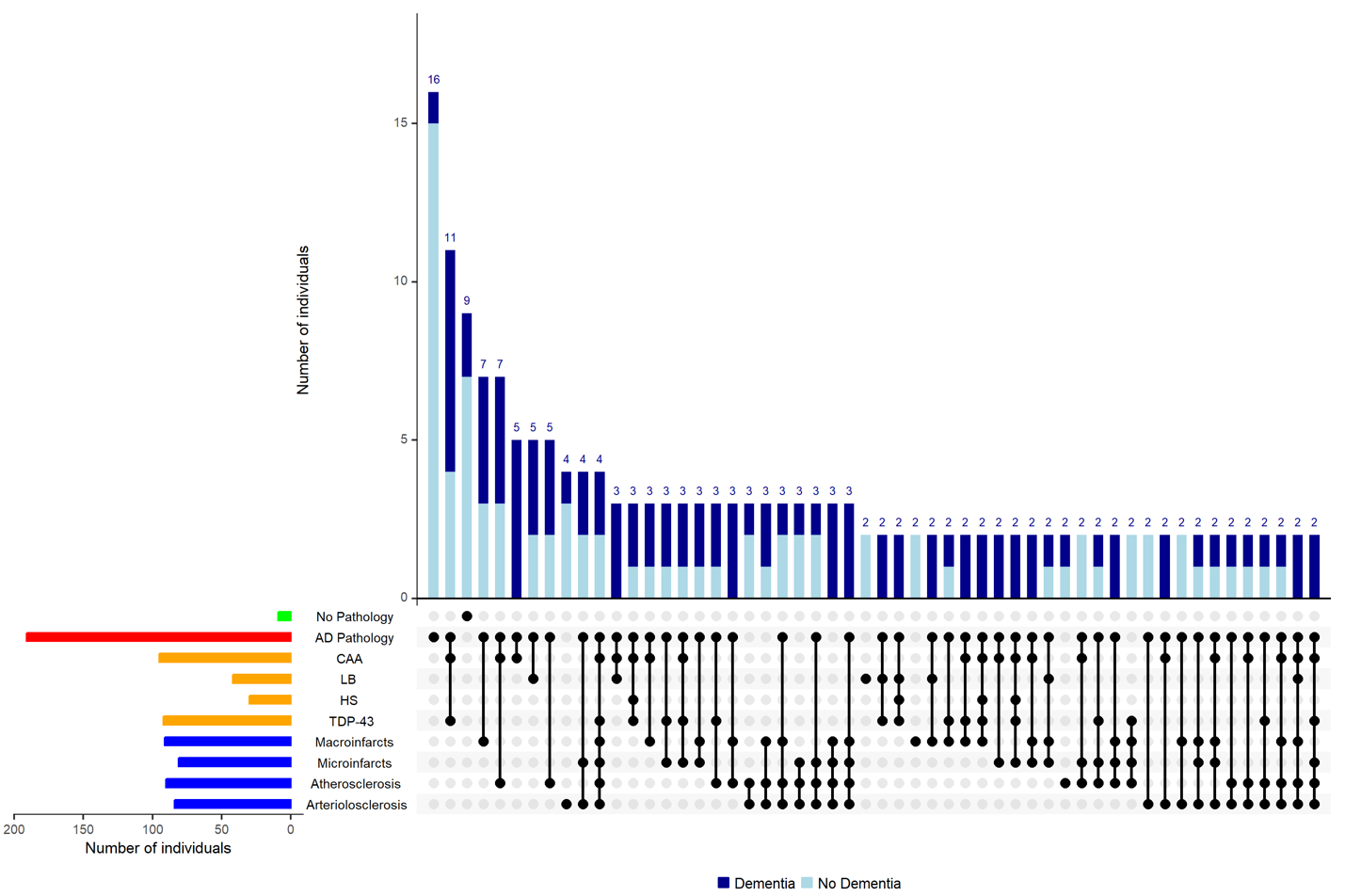

**S. Figure 2.** **UpSet figure of the distribution and frequency of mixed pathologies for subtype N_1_.** The bar chart in the lower left corner shows the frequencies of individual neuropathologic indices investigated in this study. Connected black dots on the x-axis indicate the specific combination of neuropathology represented (we show only combinations of 2+ pathologies). Histograms in the main panel show the frequencies of the neuropathologic indices for participants with and without dementia proximate to death, ordered by their frequency. The height of each bar corresponds to the number of persons with each combination. As illustrated in the figure, neuropathologic indices frequently co-occur and the most common at pathologic AD, followed by pathologic AD and a neurodegenerative pathology, most commonly, cerebral amyloid angiopathy. AD=Alzheimer’s disease. CAA= cerebral amyloid angiopathy; LB=Lewy bodies. HS=hippocampal sclerosis. TDP-43=transactivate response DNA-binding protein 43.

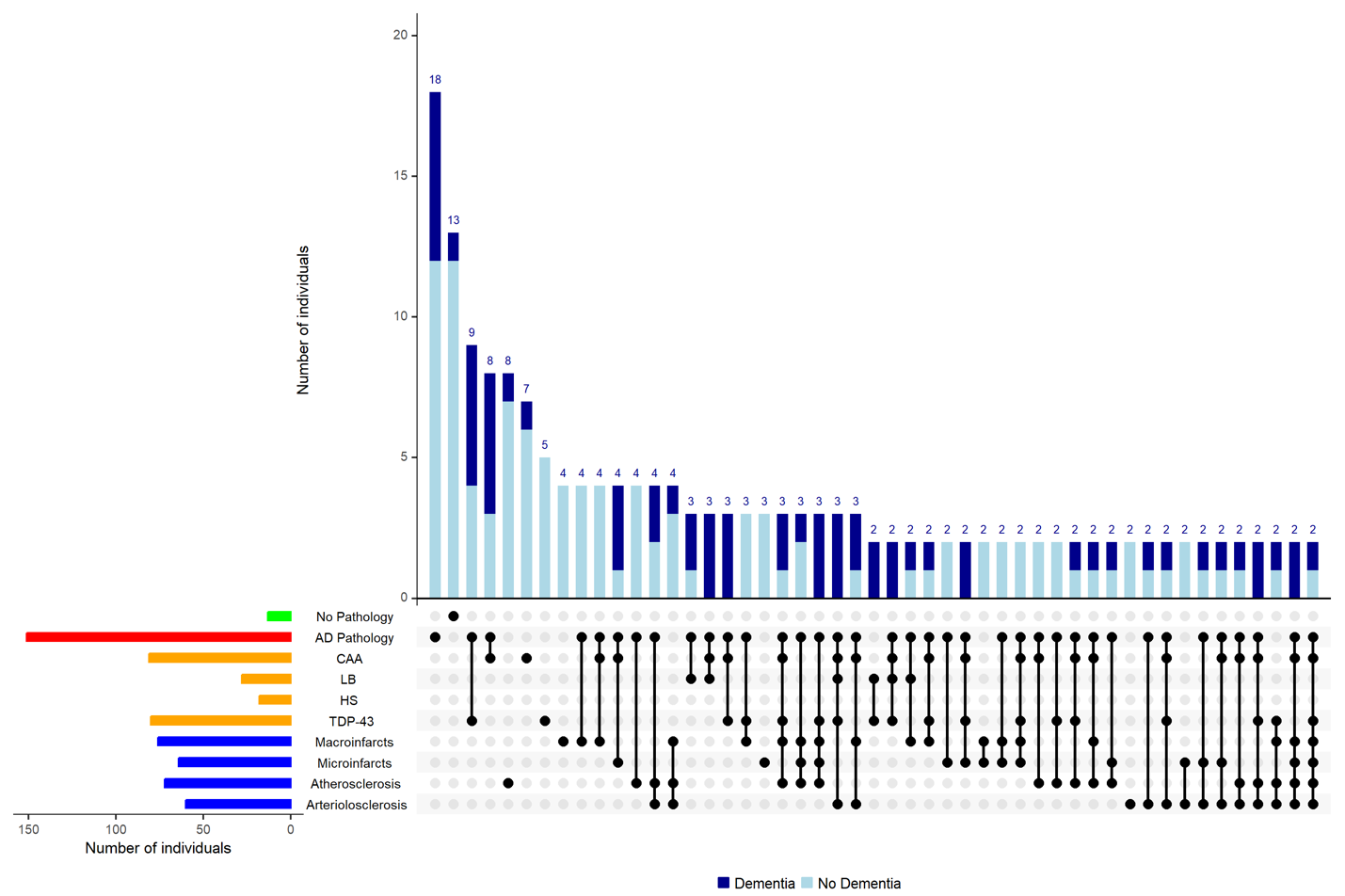

**S. Figure 3.** **UpSet figure of the distribution and frequency of mixed pathologies for subtype N_2_.** The bar chart in the lower left corner shows the frequencies of individual neuropathologic indices investigated in this study. Connected black dots on the x-axis indicate the specific combination of neuropathology represented (we show only combinations of 2+ pathologies). Histograms in the main panel show the frequencies of the neuropathologic indices for participants with and without dementia proximate to death, ordered by their frequency. The height of each bar corresponds to the number of persons with each combination. As illustrated in the figure, neuropathologic indices frequently co-occur and the most common at pathologic AD, followed by pathologic AD and a neurodegenerative pathology, most commonly, cerebral amyloid angiopathy. AD=Alzheimer’s disease. CAA= cerebral amyloid angiopathy; LB=Lewy bodies. HS=hippocampal sclerosis. TDP-43=transactivate response DNA-binding protein 43.

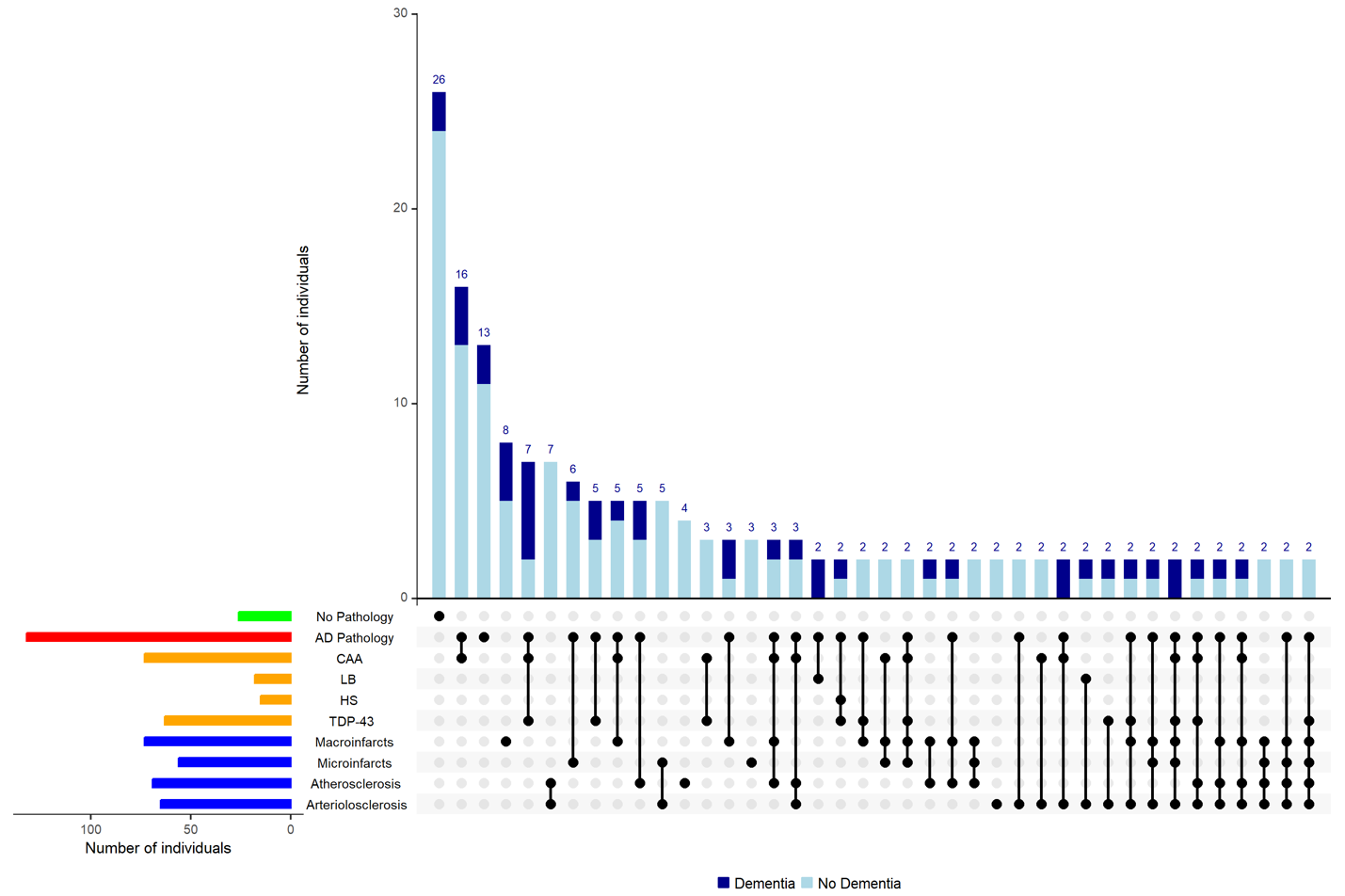

**S. Figure 4.** **UpSet figure of the distribution and frequency of mixed pathologies for subtype N_3_.** The bar chart in the lower left corner shows the frequencies of individual neuropathologic indices investigated in this study. Connected black dots on the x-axis indicate the specific combination of neuropathology represented (we show only combinations of 2+ pathologies). Histograms in the main panel show the frequencies of the neuropathologic indices for participants with and without dementia proximate to death, ordered by their frequency. The height of each bar corresponds to the number of persons with each combination. As illustrated in the figure, neuropathologic indices frequently co-occur and the most common at pathologic AD, followed by pathologic AD and a neurodegenerative pathology, most commonly, cerebral amyloid angiopathy. AD=Alzheimer’s disease. CAA= cerebral amyloid angiopathy; LB=Lewy bodies. HS=hippocampal sclerosis. TDP-43=transactivate response DNA-binding protein 43.

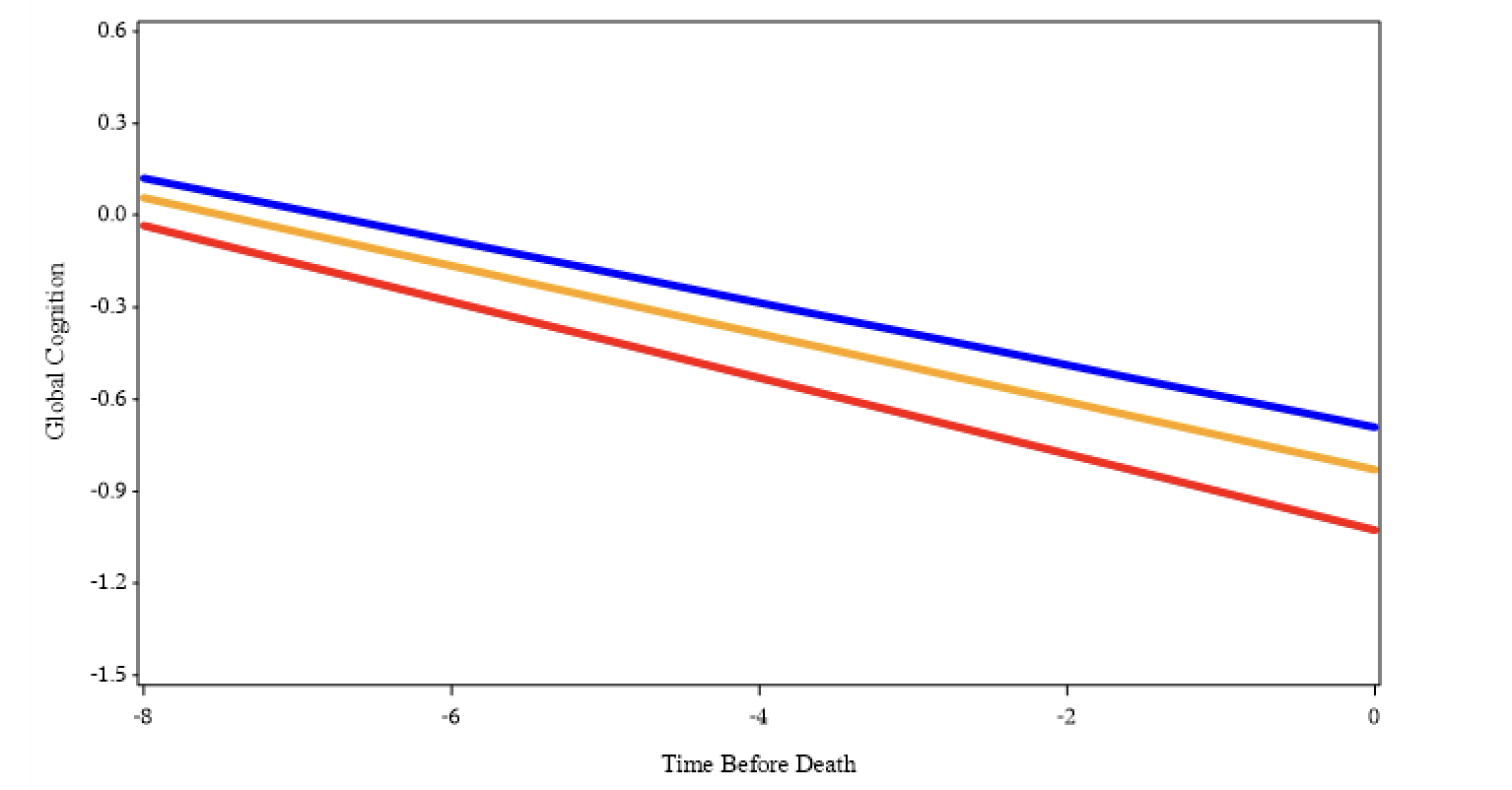

**S. Figure 5.** Predicted trajectories in global cognition during the last eight years of life for individuals in 10^th^ (blue line), 50^th^ (orange line), and 90^th^ (red line) percentile of pseudotime for an 89-year-old female with 16 years of education. Compared to the participant in the 50^th^ percentile, decline was 12% faster for the participant in the 90^th^ percentile and 9% slower for the participant in the 10^th^ percentile.
